# How does the magnitude of genetic variation affect eco-evolutionary dynamics in character displacement?

**DOI:** 10.1101/2021.02.26.432976

**Authors:** Keiichi Morita, Masato Yamamichi

## Abstract

While previous studies on character displacement tended to focus on trait divergence and convergence as a result of long-term evolution, recent studies suggest that character displacement can be a special case of evolutionary rescue, where rapid evolution prevents population extinction by weakening negative interspecific interactions. When the magnitude of genetic variation is small, however, the speed of trait divergence can be slow and populations may go extinct before the completion of character displacement. Here we analyzed a simple model to examine how the magnitude of genetic variation affects evolutionary rescue via ecological and reproductive character displacement that weakens resource competition and reproductive interference, respectively. We found that the large additive genetic variance is more important for preventing extinction in reproductive character displacement than in ecological character displacement. This is because reproductive interference produces a locally stable coexistence equilibrium with positive frequency-dependence (i.e., minority disadvantage) whereas ecological character displacement results in a globally stable coexistence equilibrium. Furthermore, population extinction becomes less likely when ecological and reproductive character displacement occur simultaneously due to positive covariance between ecological and reproductive traits. Our results suggest that while reproductive character displacement may be rarer than ecological character displacement, it is more likely to occur when there exists positive trait covariance, such as the case of a magic trait in reinforcement of speciation processes.

## Introduction

Character displacement, in which sympatric populations show trait divergence for reducing negative interspecific interactions (Brown and Wilson 1956, Grant 1972, Pfennig and Pfennig 2009, Pfennig and Pfennig 2020), is a classical topic in ecology and evolutionary biology for understanding resource partitioning, species coexistence, reinforcement, and speciation (Schluter 2000, Dayan and Simberloff 2005, Stuart and Losos 2013, Germain et al. 2018). While previous studies tended to focus on trait divergence and convergence as an outcome of long-term evolution (Lawlor and Maynard Smith 1976, Slatkin 1980, Taper and Case 1985, Abrams 1986, Doebeli 1996, Goldberg and Lande 2006, Konuma and Chiba 2007, Abrams and Cortez 2015), recent studies suggest that character displacement can be driven by rapid evolution (Grant and Grant 2006, Stuart and Losos 2013) and be a special case of evolutionary rescue (Bell 2017), where rapid evolution prevents population extinction by weakening negative interspecific interactions (Yamaguchi and Iwasa 2013, Kyogoku and Wheatcroft 2020). These days it is increasingly becoming clear that adaptive evolution can be rapid enough to affect various ecological dynamics (Hairston et al. 2005, Schoener 2011, Hendry 2016), and hence it is crucial to consider eco-evolutionary dynamics of character displacement where ecological and evolutionary processes are dynamically interacting on the same timescales (Dayan and Simberloff 2005, Stuart and Losos 2013, Kyogoku and Wheatcroft 2020).

Previous studies demonstrated that the amount of intraspecific genetic variation is an important parameter in eco-evolutionary dynamics: increasing standing genetic variation promotes evolutionary rescue, where adaptive evolution to an altered environment prevents population extinction (Gomulkiewicz and Holt 1995, Agashe 2009, Bell 2017), promotes coexistence of competing species (Vasseur et al. 2011, Mougi 2013, Yamamichi et al. 2020, Yamamichi and Letten 2021), and can stabilize or destabilize predator-prey population cycles (Abrams and Matsuda 1997, Becks et al. 2010, Mougi and Iwasa 2010, Cortez 2016, Cortez and Patel 2017, Cortez 2018). We propose that the magnitude of genetic variation (and the resultant speed of evolution) is also an important parameter in character displacement. This is because, if trait divergence is slow, populations may go extinct before the completion of character displacement due to the negative interspecific interactions. Traditionally researchers have studied two types of negative interspecific interactions in character displacement (Pfennig and Pfennig 2020): resource competition, a negative interspecific interaction through exploitation of shared resources, and reproductive interference, a negative interspecific interaction in the mating process caused by incomplete species recognition (Gröning and Hochkirch 2008). Ecological and reproductive character displacement weakens resource competition and reproductive interference, respectively (Goldberg and Lande 2006, Pfennig and Pfennig 2020). Previous studies on eco-evolutionary dynamics often focused on ecological (trophic) traits (Hairston et al. 2005, Schoener 2011, Hendry 2016), but recent studies emphasized the importance of sexual traits (Giery and Layman 2019, Svensson 2019, Yamamichi et al. 2020). In this study, we ask how the magnitude of genetic variation (i.e., the speed of evolution) affects evolutionary rescue via ecological and reproductive character displacement.

We developed a simple model of character displacement by adding evolutionary dynamics of quantitative traits (Iwasa et al. 1991) to a model with resource competition and reproductive interference (Schreiber et al. 2019). We conducted numerical simulations of the minimum model and found that additive genetic variance of the reproductive trait (e.g., flower colors) in reproductive character displacement is more crucial for preventing extinction than that of the ecological trait (e.g., root depths) in ecological character displacement. This is because reproductive interference produces positive frequency-dependence in community dynamics (Kuno 1992, Yoshimura and Clark 1994, Kishi and Nakazawa 2013, Schreiber et al. 2019) and trait divergence needs to occur before population decline. Furthermore, extinction becomes less likely when ecological and reproductive character displacement occur simultaneously by a positive genetic covariance between ecological and reproductive traits (Konuma and Chiba 2007, Kyogoku and Kokko 2020) as like a magic trait, a trait subject to divergent selection and contributing to assortative mating, promotes speciation (Servedio et al. 2011). Our results suggest that reproductive character displacement may be rarer than ecological character displacement, but it may be promoted by ecological trait divergence.

## Models

We consider a discrete-time model of two populations with densities *N*_1_ and *N*_2_ interacting with resource competition and reproductive interference (Schreiber et al. 2019):

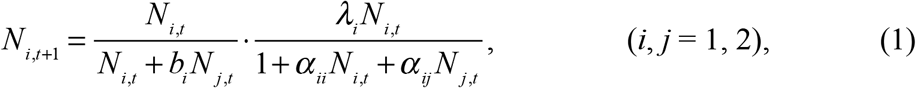

where *λ*_*i*_ is the maximum per-capita fecundity, *b*_*i*_ is the strength of reproductive interference, and *α*_*ii*_ and *α*_*ij*_ are the strength of intraspecific and interspecific resource competition, respectively.

When the parameter *b*_*i*_ is zero, it is the classical Leslie-Gower model (Leslie and Gower 1958) that has been used to describe competition dynamics in annual plants and insects (Leslie and Gower 1958, Chesson 1994, Adler et al. 2007, Godoy and Levine 2014). Its dynamics serve as a discrete-time analogs of the classical continuous-time Lotka-Volterra competition model (Cushing et al. 2004). When *b*_*i*_ is positive, community dynamics shows positive frequency-dependence where rare species are disadvantaged relative to common species (Kuno 1992, Yoshimura and Clark 1994, Kishi and Nakazawa 2013, Schreiber et al. 2019). Therefore, even with negative frequency-dependence due to resource partitioning, rare species may go extinct. Ecological dynamics of the model (Equation (1)) were investigated in detail by the previous study (Schreiber et al. 2019), which showed that coexistence is possible when niche overlap,

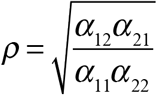

(Godoy and Levine 2014), and the intensity of reproductive interference, *b*_*i*_, are small (the orange region in Figure 1a).

**Figure 1.**
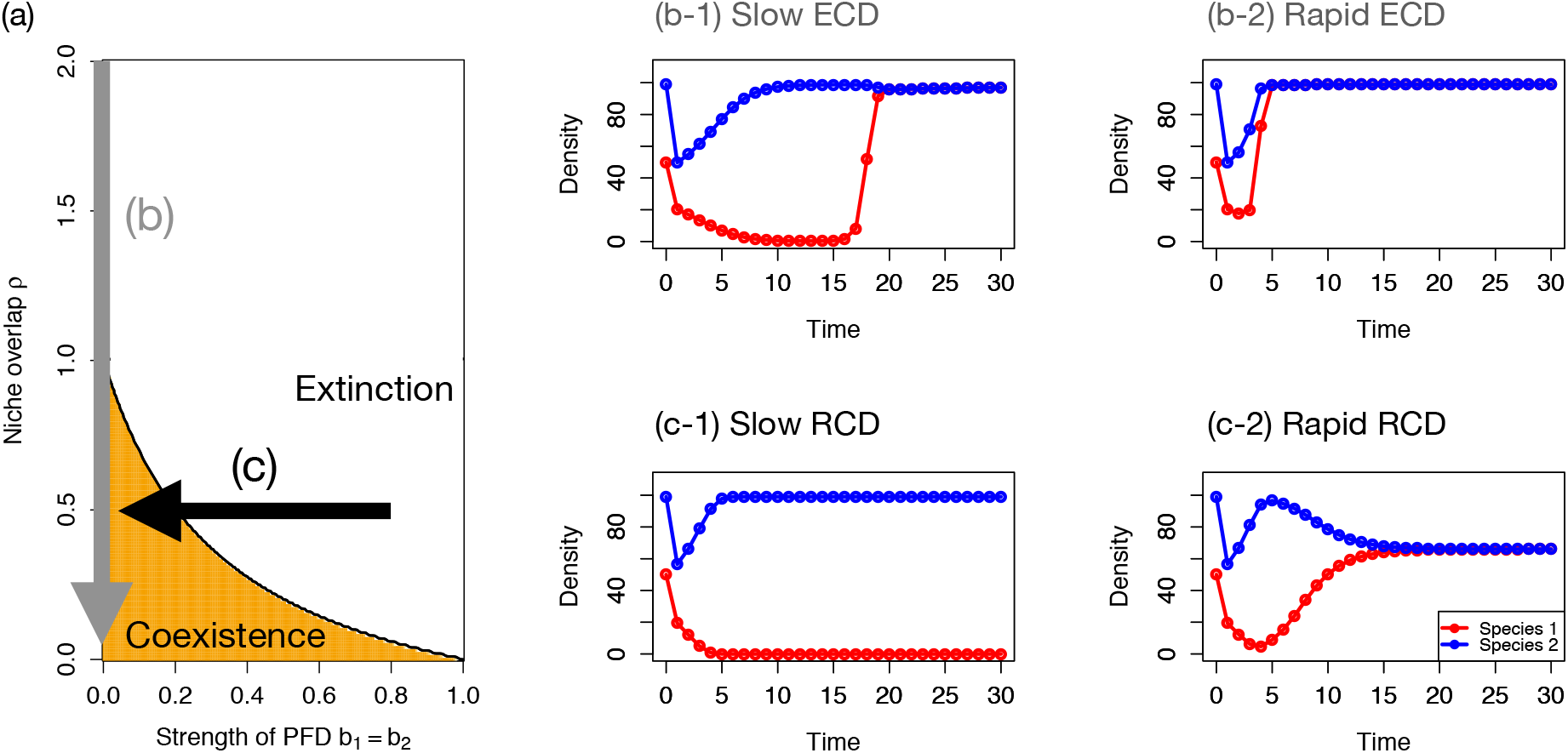
(a) Coexistence conditions in the ecological model (Equation (1)) (Schreiber et al. 2019). Coexistence is possible in the orange region where niche overlap is small and reproductive interference is weak. The gray and black arrows represent Ecological Character Displacement (ECD) and Reproductive Character Displacement (RCD), respectively. Parameter values are *α*_11_ = *α*_22_ = 1 and *λ*_1_ = *λ*_2_ = 100 and we assumed *α*_12_ = *α*_21_. (b) ECD results in coexistence irrespective of the magnitude of genetic variation (b-1: *G*_*x*_ = 0.1, b-2: *G*_*x*_ = 1). The initial density is *N*_1,0_ = 50 and parameter values are *α*_12,max_ = *α*_21,max_ = 2 and *b*_1,max_ = *b*_2,max_ = 0. (c) RCD promotes coexistence with rapid evolution due to large genetic variance (c-2: *G*_*y*_ = 1) whereas it results in extinction of rare species with small genetic variance (c-1: *G*_*y*_ = 0.1). Here parameter values are *α*_12,max_ = *α*_21,max_ = 0.5 and *b*_1,max_ = *b*_2,max_ = 0.8.

We assume the species *i* have two quantitative traits, *x*_*i*_ and *y*_*i*_, that affect the strength of interspecific resource competition (*α*_*ij*_) and reproductive interference (*b*_*i*_), respectively. We also assume that the difference between the two species determines the interaction strengths as follows:

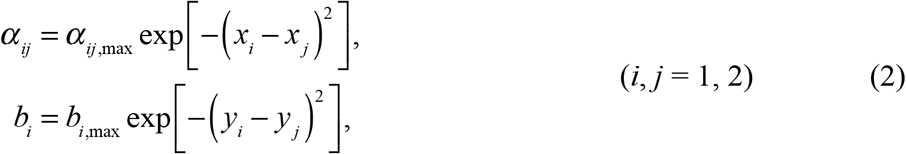

where *α*_*ij*,max_ and *b*_*i*,max_ are maximum strengths of interspecific resource competition and reproductive interference, respectively.

In character displacement, selection increases the difference between trait values so that negative interspecific interactions become weaker. Adaptive evolution of the quantitative traits occurs along the fitness gradient as follows (Iwasa et al. 1991):

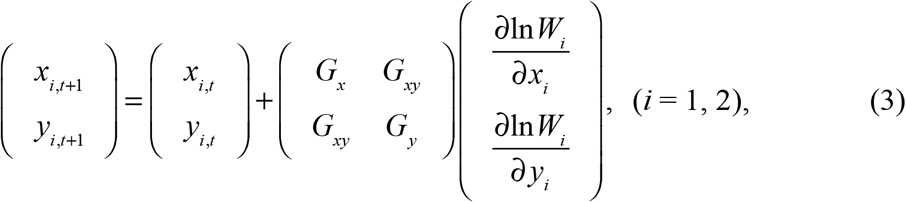

where *W*_*i*_ = *N*_*i,t*+1_/*N*_*i,t*_ is fitness, *G*_*x*_ and *G*_*y*_ are additive genetic variance of the reproductive and ecological traits, respectively, and *G*_*xy*_ is additive genetic covariance. We assume that many loci affect the traits additively, and that the distribution of the trait in the population concentrates sharply around the population average (but does not need to be normal) (Iwasa et al. 1991). Positive (negative) covariance means evolution of a trait promotes (hinder) evolution of the other trait. The speed of evolution is determined by genetic (co)variance and the fitness gradient. Because the fitness gradient depends on population densities and trait values, we can control the speed of evolution by changing values of genetic (co)variance. We assume that fitness gradient is only affected by interspecific interactions that prefer the larger difference in trait values. Thus, there is no internal stable equilibrium of the trait values and they may keep evolving to positive or negative infinity.

We consider three scenarios: (1) ecological character displacement, where there is no reproductive interference (*b*_*i*_ = 0) and ecological trait divergence weakens interspecific resource competition (the gray arrow in Figure 1a), (2) reproductive character displacement, where there is ecological niche differentiation (*ρ* < 1) and reproductive trait divergence weakens reproductive interference (the black arrow in Figure 1a), and (3) simultaneous ecological and reproductive character displacement, where two traits diverge simultaneously. We compare the effects of genetic variance on the scenarios (1) and (2), and then examine the effects of positive and negative genetic covariance in the scenario (3). When a population of species 1 migrates to a habitat where a population of species 2 dominates, rapid evolution of species 1 may allow it to avoid its extinction (i.e., evolutionary rescue) through trait divergence. Although the small population size of migrating species 1 may result in low genetic variation and hinder evolutionary responses, recent studies revealed the prevalence of rapid adaptive evolution of invasive species (Prentis et al. 2008). For simplicity, we assumed that trait values of species 2 are constant (*x*_2_ = *y*_2_ = 0) and its initial density is at a carrying capacity (i.e., *N*_2,0_ = (*λ*_2_ – 1)/*α*_22_). The initial trait values of species 1 are assumed to be slightly larger than the fixed trait values of species 2 (*x*_1,0_ = *y*_1,0_ = 0.1), so selection on species 1 due to interspecific interactions drives trait values to positive infinity. Then we changed additive genetic (co)variance of species 1 to explore the effect of the speed of evolution on eco-evolutionary dynamics of character displacement. We employed the basic parameter values of Schreiber et al. (2019) and used R for simulations (R Core Team 2019).

## Results

Ecological character displacement decreases niche overlap (the gray arrow in Figure 1a) and promotes coexistence of two species even with small genetic variance (Figure 1b). On the other hand, reproductive character displacement decreases the strength of reproductive interference (the black arrow in Figure 1a) but results in deterministic extinction of initially rare species when genetic variance is small and evolution is slow (Figure 1c). The difference in the importance of genetic variance arises due to the presence of positive frequency-dependence. Even in the coexistence region in Figure 1a, coexistence is possible with a limited set of initial conditions with reproductive interference (Schreiber et al. 2019).

This point can be clearly shown by using nullclines analyses. Before character displacement, strong niche overlap in resource competition and reproductive interference results in extinction of rare species (i.e., positive frequency-dependence: Figure 2a-1, 2b-1). Character displacement produces a stable coexistence equilibrium, but rare species still go extinct when there is reproductive interference (Figure 2a-2, 2b-2) (Schreiber et al. 2019). The difference arises because ecological character displacement produces a globally stable coexistence equilibrium (Figure 2a-2), whereas reproductive character displacement produces a locally stable coexistence equilibrium (Figure 2b-2). Even after divergence of the reproductive trait, when the densities of the two species are outside of the basin of attraction to the coexistence equilibrium, one of the two species goes extinct deterministically (Figure 2b-2).

**Figure 2.**
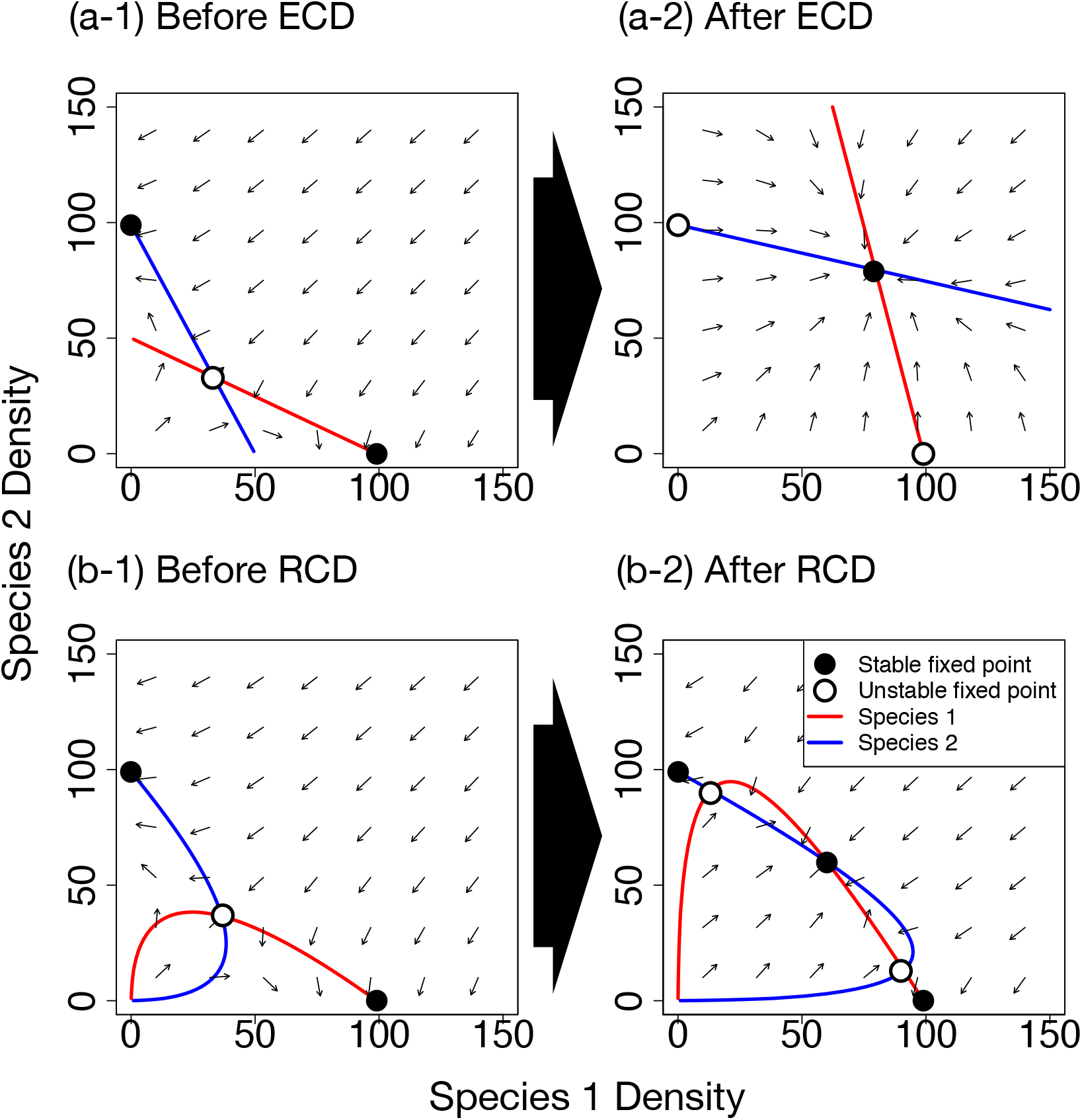
Nullclines before and after character displacement. (a) Ecological Character Displacement (ECD) changes the dynamics from that with two locally stable extinction equilibria to that with a globally stable coexistence equilibrium. (b) Reproductive Character Displacement (RCD) changes the dynamics from that with two locally stable extinction equilibria to that with a locally stable coexistence equilibrium. Even after RCD, coexistence does not occur when the density of one species is very small. Here *x*_1_ = *y*_1_ = 0.1 in a-1 and b-1, *x*_1_ = *y*_1_ = 1.45 in a-2 and b-2, and other parameter values are the same as Figure 1.

As we defined that trait differences weaken interspecific resource competition and reproductive interference (Equation (2)), bifurcations occur as we increase trait differences. In ecological character displacement, the bifurcation produces a globally stable coexistence equilibrium (Figure 3a), whereas the bifurcation in reproductive character displacement produces a locally stable coexistence equilibrium with two unstable equilibria (Figure 3b). If evolution is slow due to small additive genetic variance, rare species cannot cross the unstable equilibrium, and attracted to the deterministic extinction equilibrium (Figure 3b-1); on the other hand, rapid evolution makes it possible for rare species to cross the unstable density equilibrium to the basin of attraction toward the locally stable coexistence equilibrium (Figure 3b-2).

**Figure 3.**
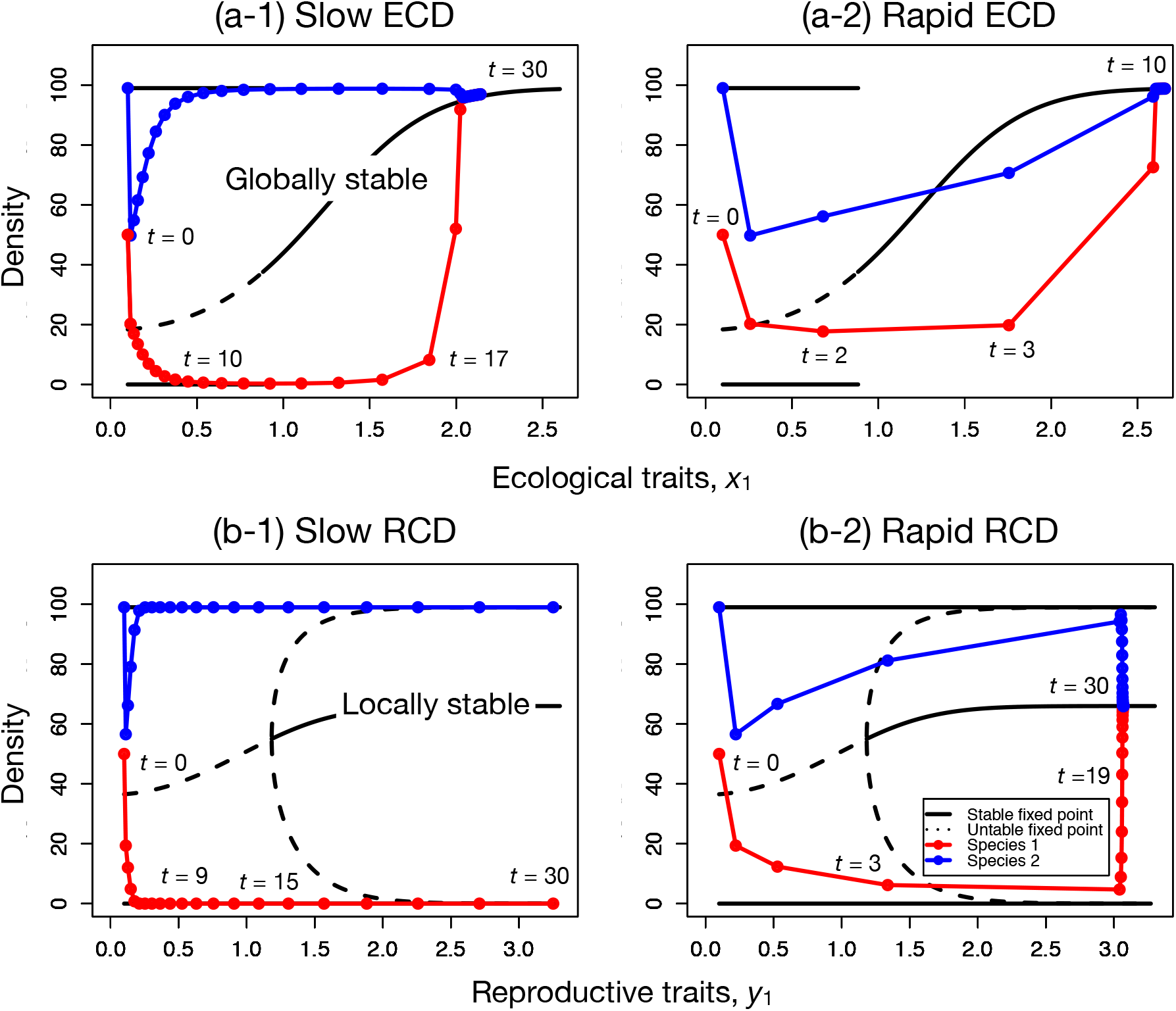
Bifurcation plots along ecological and reproductive trait values. Dashed and solid lines indicate unstable and stable equilibria, respectively. Blue and red solid lines represent simulation trajectories of species 1 and 2, respectively. (a) Ecological Character Displacement (ECD) produces a globally stable coexistence equilibrium. (b) Reproductive Character Displacement (RCD) produces a locally stable coexistence equilibrium and two unstable equilibria. Parameter conditions are the same as Figure 1.

The phase diagram of initial trait difference versus additive genetic variance shows that ecological character displacement results in coexistence irrespective of parameter values (Figure 4a). When the two parameters are small, transient dynamics can become very long, but the system is eventually attracted to the coexistence equilibrium as we did not consider demographic stochasticity. On the other hand, reproductive character displacement requires large additive genetic variance (and the resultant rapid evolution) as well as large initial trait difference for coexistence (Figure 4b).

**Figure 4.**
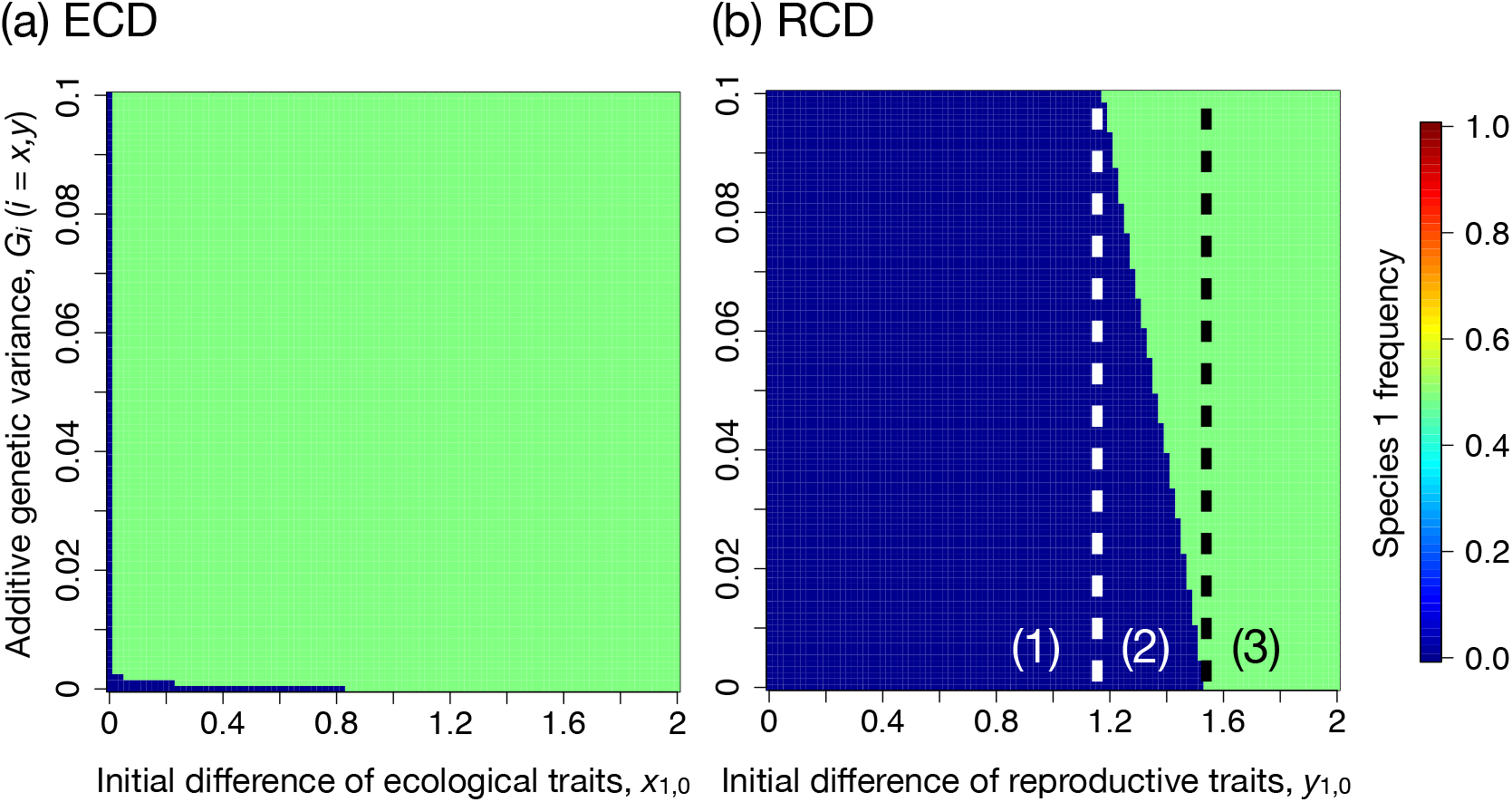
Phase diagrams showing the effects of initial trait differences (*X*-axis) and additive genetic variance (*Y*-axis) on the frequency of species 1 in the community, *N*_1_/(*N*_1_ + *N*_2_), after (a) 1000 and (b) 50 time-steps. (a) Ecological Character Displacement (ECD) almost always results in coexistence. (b) Reproductive Character Displacement (RCD) promotes coexistence with the large initial trait differences and additive genetic variance. In region (1), the rare species goes extinct irrespective of additive genetic variance. In region (3), coexistence is possible. In region (2), coexistence depends on additive genetic variance. The initial density is *N*_1,0_ = 10, and other parameter values are the same as Figure 1.

We further consider the situation where two types of character displacement occur simultaneously and are linked by genetic covariance between ecological and reproductive traits, *G*_*xy*_, of Equation (3). With positive genetic covariance, coexistence becomes more likely because the two types of displacement facilitate each other (Figure 5). However, when genetic covariance is negative, the divergence of ecological traits can prevent the divergence of reproductive traits and vice versa, which may prevent the promotion of coexistence (Figure 5).

**Figure 5.**
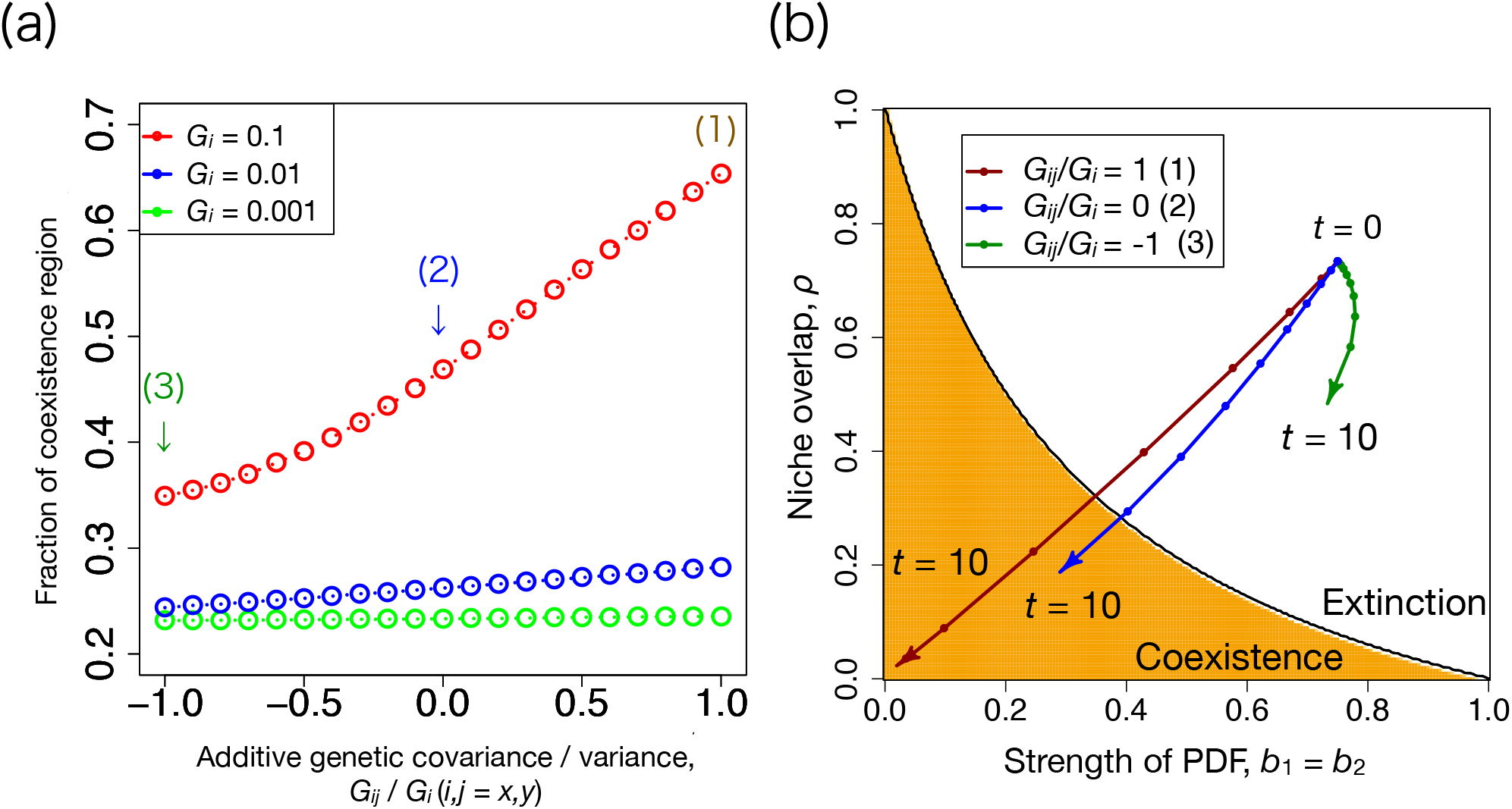
(a) Positive additive genetic covariance promotes coexistence. We show three situations where additive genetic variance, *G*_*x*_ = *G*_*y*_ = 0.001 (green), 0.01 (blue), and 0.1 (red). *X*-axis is the ratio of genetic covariance to additive genetic variance (*G*_*xy*_/*G*_*i*_, *i* = *x, y*), and *Y*-axis is the fraction of coexistence region in phase diagrams as Figure 4. The initial density is *N*_1,0_ = 10, parameter values are *b*_1,max_ = *b*_2,max_ = 0.5 and *α*_12,max_ = *α*_21,max_ = 3, and other parameter values are the same as Figure 1. (b) Rapid evolution promotes trait divergence and changes the parameter condition from the extinction (white) region to the coexistence (orange) region. We show three situations where (1) additive genetic variance and covariance are the same, *G*_*x*_ = *G*_*y*_ = *G*_*xy*_ = 0.1 (brown), (2) genetic covariance is zero (blue), and (3) genetic covariance is negative, *G*_*xy*_ = −0.1 (green).

With the negative genetic covariance, it is even possible for reproductive interference to temporarily become stronger than the initial value through adaptive evolution. This occurs because evolution of the reproductive trait is affected by evolution of the ecological trait weakening resource competition (the green points initially move toward bottom-right in Figure 5b). However, when the evolving reproductive trait of species 1 becomes smaller than the fixed reproductive trait of species 2 (*y*_1,*t*_ < *y*_2_ = 0), the two types of character displacement are facilitated by the negative genetic covariance. Beyond this point, increasing the ecological trait value and decreasing the reproductive trait value weaken resource competition and reproductive interference simultaneously (the green points eventually move toward bottom-left in Figure 5b). This occurs because we assumed that the absolute difference of trait values determines interaction strength (Equation (2)). It is also possible for resource competition to be temporarily intensified by weakening reproductive interference (a trajectory moves to top-left in the phase diagram from the initial point in Figure 5b) due to the negative genetic covariance and smaller *α*_*ij*,max_ (data not shown).

## Discussion

By simulating eco-evolutionary models, we revealed that rapid evolution can prevent extinction of initially rare species by ecological and reproductive character displacement. Therefore, character displacement can be a special case of evolutionary rescue, where adaptation to interspecific interactions prevents extinction (Yamaguchi and Iwasa 2013, Bell 2017, Kyogoku and Wheatcroft 2020). In addition, we showed that the magnitude of genetic variance is more important in reproductive character displacement due to positive frequency-dependence (Figures 1-4). While previous studies on character displacement and species coexistence tended to focus on competitive exclusion, recent studies have revealed that reproductive interference is an important factor that promotes extinction of rare species via positive frequency-dependence (i.e., reproductive exclusion: (Pfennig and Pfennig 2009)) by empirical (Kawatsu and Kishi 2018, Christie and Strauss 2020) and theoretical (Kuno 1992, Yoshimura and Clark 1994, Kishi and Nakazawa 2013, Schreiber et al. 2019) approaches. Reproductive character displacement produces a locally stable coexistence equilibrium with minority disadvantage (Fig. 2b, 3b), but ecological character displacement produces a globally stable coexistence equilibrium (Fig. 2a, 3a). This difference indicates that reproductive character displacement may not occur as often as ecological character displacement in the wild due to the lack of genetic variation. Distribution of two closely related species may overlap and there may be character displacement if the two species compete for shared resources, but the distribution overlap may be rare if the two species interact via reproductive interference. However, it should be noted that our model did not consider extinction due to demographic stochasticity, and stochastic extinction may occur due to slow evolution even in ecological character displacement.

Our results further indicate that reproductive character displacement can be promoted by ecological character displacement when there is a positive trait covariance as previous studies pointed out (Konuma and Chiba 2007, Kyogoku and Kokko 2020). This is similar to the situation of magic traits in the context of speciation processes, where ecological adaptation and assortative mating are realized by a single trait or some genetic linkages (Servedio et al. 2011). Actually, reproductive character displacement is similar to reinforcement in the speciation processes (Pfennig and Pfennig 2009), and Liou and Price (1994) theoretically showed that the small initial trait difference can result in extinction (as our Figure 4b) during the process of speciation by reinforcement. However, they also showed that extinction can be avoided by producing a hybrid swarm when hybrid fitness is high. While our model assumes that there are only two outcomes in reproductive character displacement (i.e., extinction or coexistence after the completion of displacement), future studies should consider the three possibilities (i.e., extinction, coexistence, or hybridization) for fully understanding the eco-evolutionary dynamics of speciation by reinforcement/reproductive character displacement.

Finally, we showed that negative covariance of the ecological and reproductive traits can produce a transient increase in one of the negative interspecific interactions (Figure 5b). This indicates the difficulty of understanding evolutionary dynamics by focusing on a single trait (Kopp and Matuszewski 2014, Yamamichi et al. 2019). For example, when an alien plant species invades certain habitats, subsequent evolution that strengthens reproductive interference (e.g., convergence of flower colors) may also weaken resource competition (e.g., divergence of root depths) when there is a genetic linkage between the two traits.

Recent studies on evolutionary rescue have explored the effects of competitive (Osmond and de Mazancourt 2013), predator-prey (Yamamichi and Miner 2015), and mutualistic (Jones and Gomulkiewicz 2012) interactions on population extinction in changing environments. On the other hand, our study examined how adaptation to negative interspecific interactions prevent extinction in a constant environment. It will be important to integrate theory of character displacement, evolutionary rescue, and species coexistence through the lens of eco-evolutionary dynamics for better understanding the maintenance of biodiversity in future studies.

In conclusion, we showed that reproductive trait evolution needs to be rapid in order to prevent extinction of rare species, and that positive genetic covariance between ecological and reproductive traits contributes to species coexistence. Our simple and phenomenological model of character displacement is different from more mechanistic models in previous studies with a single trait axis for divergence (Slatkin 1980, Taper and Case 1985, Goldberg and Lande 2006, Konuma and Chiba 2007) and explicit resource dynamics (Lawlor and Maynard Smith 1976, Abrams 1986, Abrams and Cortez 2015), and will be useful for linking character displacement and coexistence theory (Germain et al. 2018). It should be noted that our trait dynamics are affected only by interspecific interactions, and trait dynamics do not show trait convergence (Grant 1972). Future studies should add stabilizing selection through a trade-off between interspecific competition and intraspecific interaction (Vasseur et al. 2011, Mougi 2013) or fecundity. We assumed that the resident species 2 does not evolve for simplicity. This may be justified, at least at the initial transient stage of character displacement, because selection pressure from an initially rare species 1 may be weak. However, species 2 may have higher genetic variation due to its large population size, and coevolution may make coexistence easier. Following the approach by previous studies in quantitative genetics (Iwasa et al. 1991) and recent studies on eco-evolutionary dynamics (Gomulkiewicz and Holt 1995, Abrams and Matsuda 1997, Mougi and Iwasa 2010, Vasseur et al. 2011, Mougi 2013, Cortez 2016, Cortez and Patel 2017, Cortez 2018, Yamamichi et al. 2020, Yamamichi and Letten 2021), we assumed constant additive genetic variance. It should be noted that our simplifying assumption of constant variance is unlikely to hold in nature. Displacement in itself dynamically changes additive genetic variance (Slatkin 1980, Taper and Case 1985, Doebeli 1996). Also, rare species are likely to have small additive genetic variance which may hinder their capacity for rapid evolution (Prentis et al. 2008, Bell 2017). Therefore, future studies may benefit from using more genetically explicit modeling (Doebeli 1996) or individual-based simulations. Such simulations will be useful for adding further complexities including spatial structure with migration from neighboring habitats, hybridization, coevolution, resource dynamics, and demographic stochasticity.

## Acknowledgements

We thank S. P. Hart for helpful discussion and S. Dobata, A. R. M. James, K. Lyberger, Y. Okuzaki, and N. Shinohara for helpful comments. MY was supported by the Japan Society for the Promotion of Science (JSPS) Grant-in-Aid for Scientific Research (KAKENHI) 16K18618, 18H02509, 19K16223, and 20KK0169.

